# ACE2 binding is an ancestral and evolvable trait of sarbecoviruses

**DOI:** 10.1101/2021.07.17.452804

**Authors:** Tyler N. Starr, Samantha K. Zepeda, Alexandra C. Walls, Allison J. Greaney, David Veesler, Jesse D. Bloom

**Author notes:** These authors contributed equally (Tyler N. Starr, Samantha K. Zepeda).

## Abstract

Two different sarbecoviruses have caused major human outbreaks in the last two decades^1,2^. Both these sarbecoviruses, SARS-CoV-1 and SARS-CoV-2, engage ACE2 via the spike receptor-binding domain (RBD)^2–6^. However, binding to ACE2 orthologs from humans, bats, and other species has been observed only sporadically among the broader diversity of bat sarbecoviruses^7–11^. Here, we use high-throughput assays^12^ to trace the evolutionary history of ACE2 binding across a diverse range of sarbecoviruses and ACE2 orthologs. We find that ACE2 binding is an ancestral trait of sarbecovirus RBDs that has subsequently been lost in some clades. Furthermore, we demonstrate for the first time that bat sarbecoviruses from outside Asia can bind ACE2. In addition, ACE2 binding is highly evolvable: for many sarbecovirus RBDs there are single amino-acid mutations that enable binding to new ACE2 orthologs. However, the effects of individual mutations can differ markedly between viruses, as illustrated by the N501Y mutation which enhances human ACE2 binding affinity within several SARS-CoV-2 variants of concern^12^ but severely dampens it for SARS-CoV-1. Our results point to the deep ancestral origin and evolutionary plasticity of ACE2 binding, broadening consideration of the range of sarbecoviruses with spillover potential.

---

Both SARS-CoV-2 and SARS-CoV-1 utilize human ACE2 as their receptor^2–6^. Sampling of bats has identified multiple lineages of sarbecoviruses^13,14^ with RBDs exhibiting different ACE2 binding properties^7–11,14–22^. Prior to the emergence of SARS-CoV-2, all bat sarbecoviruses with a demonstrated ability to bind any ACE2 ortholog contained RBDs related to SARS-CoV-1 and were sampled from *Rhinolophus sinicus* and *R. affinis* bats in Yunnan province in southwest China^7,8,11,23,24^. More recently, sarbecoviruses related to SARS-CoV-2 that bind ACE2 have been found more widely across Asia and from a broader diversity of *Rhinolophus* species^2,18,25–27^. However, ACE2 binding has not been observed within a prevalent group of sarbecovirus RBDs sampled in southeast Asia (RBD “Clade 2”)^7,8,21^, nor in distantly related sarbecoviruses from Africa and Europe (RBD “Clade 3”)^7,14^ (**Figure 1a**). Therefore, it is unknown whether ACE2 binding is an ancestral trait of sarbecovirus RBDs that has been lost in some clades, or a trait that was acquired more recently^13,14^. An additional complication is that ACE2 itself is quite variable among bats, particularly in the surface recognized by sarbecoviruses^28–30^. Therefore, it is also important to understand how difficult it is for sarbecoviruses to acquire the ability to bind new ACE2 orthologs, including that of humans.

**Figure 1.**
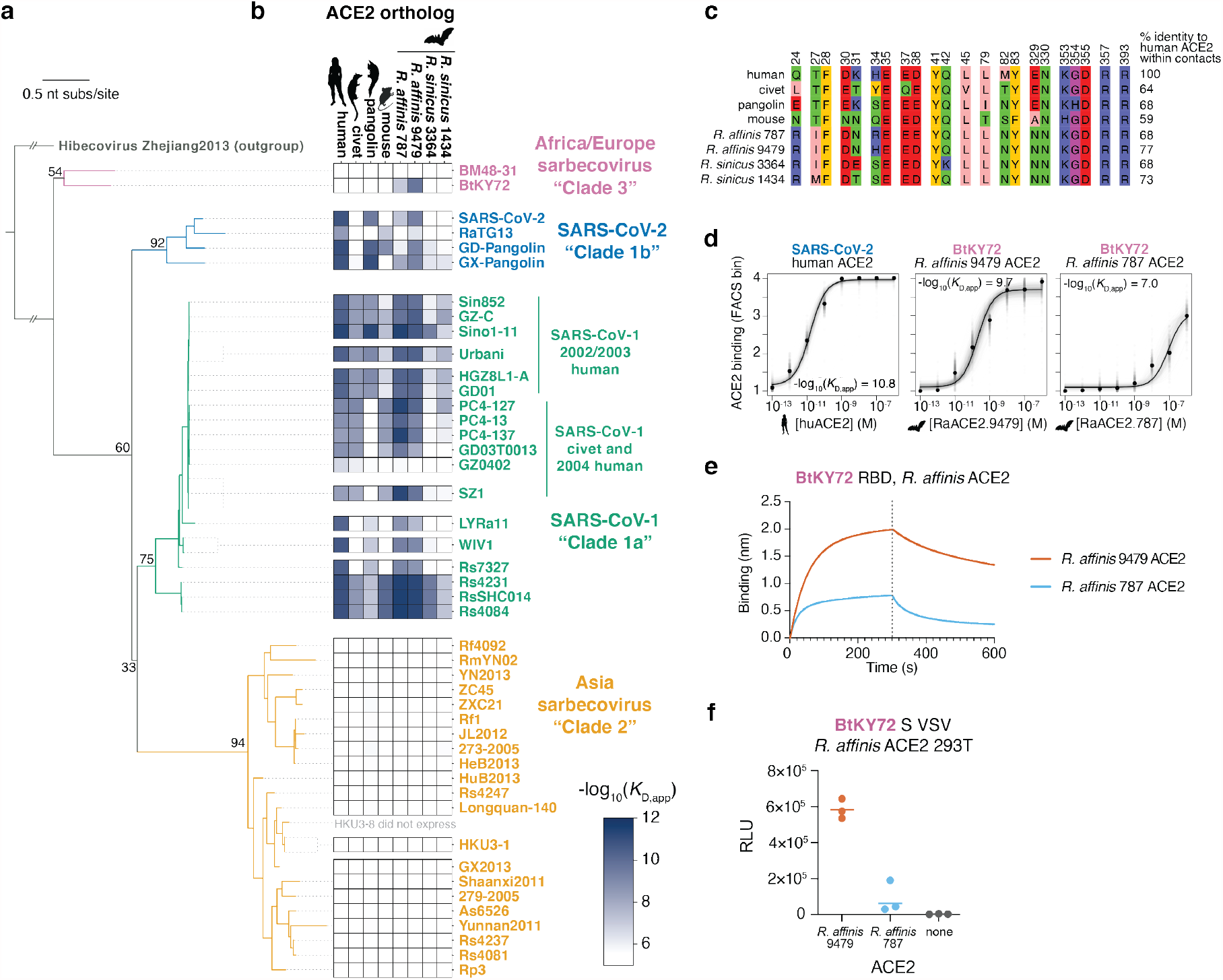
High-throughput survey of sarbecovirus ACE2 binding. **a**, Maximum likelihood phylogeny of sarbecovirus RBDs, constructed from RBD nucleotide sequences. Node labels indicate bootstrap support values. **b**, Binding avidities of sarbecovirus RBDs for eight ACE2 orthologs, determined using high-throughput yeast-displayed RBD titration assays (**Extended Data Fig. 2**). Scale bar, bottom right. **c**, Alignment of tested ACE2 orthologs within RBD-contact positions (4Å cutoff in PDB 6M0J or 2AJF). **d**, Representative binding curves from high-throughput titrations. Underlying titration curves for individual replicate-barcoded representatives of a genotype are shown in faint gray, and the average binding across all barcodes is indicated in black. **e**, Biolayer interferometry binding analysis of *R. affinis* ACE2-Fc and biotinylated BtKY72 RBD immobilized at the surface of streptavidin biosensors. Data representative of three assays using independent preparations of RBD (biological triplicate) **f**, Entry of BtKY72 spike-pseudotyped VSV particles into 293T cells transiently expressing *R. affinis* ACE2 alleles. Each point represents the mean of technical triplicates for assays performed with independent preparation of pseudoviral particles (biological replicate). Geometric mean is indicated by horizontal line. Normalized pseudovirus western blot, and mock (no S) pseudovirus entry in *R. affinis* ACE2 293T cells in **Extended Data Fig. 3a,b**.

## Complete survey of sarbecovirus ACE2 binding

To trace the evolutionary history of sarbecovirus binding to ACE2, we assembled a gene library encoding 45 sarbecovirus RBDs spanning all four major RBD phylogenetic clades (**Fig. 1a,b** and **Extended Data Fig. 1**). We cloned the RBD library into a yeast-surface display platform that enables high-throughput measurement of ACE2 binding avidities via titration assays combining fluorescence-activated cell sorting (FACS) and deep sequencing^12^ (**Extended Data Fig. 2a-d**). We also assembled a panel of recombinant, dimeric ACE2 proteins from human, civet, pangolin, mouse, and two alleles each from *R. affinis* and *R. sinicus* bats^28^ (**Fig. 1c**). The two *R. affinis* alleles represent the two distinct RBD-interface sequences found among 23 *R. affinis* bats from Yunnan and Hubei, China, whereas the two *R. sinicus* alleles represent two of the eight distinct RBD-interface sequences found among 25 *R. sinicus* bats from Yunnan, Hubei, Guangdong, and Guangxi provinces^28^. We measured the apparent dissociation constant (*K*_D,app_) of each RBD for each of the eight ACE2 orthologs (**Fig. 1b,d** and **Extended Data Fig. 2**). We performed all experiments in duplicate using independently constructed libraries, and the measurements were highly correlated between replicates (*R*^2^ > 0.99, **Extended Data Fig. 2g**).

Consistent with a prior survey of human ACE2-mediated cellular infectivity^7^, human ACE2 binding is restricted to RBDs within the SARS-CoV-1 and SARS-CoV-2 clades (**Fig. 1b**), although binding affinities vary among RBDs within these clades. Specifically, the RBDs from SARS-CoV-2 and related viruses from pangolins bind human ACE2 with high affinity, while the RBD from the bat virus RaTG13 exhibits much lower affinity^18^. The RBDs of SARS-CoV-1 isolates from the 2002-2003 epidemic bind human ACE2 strongly, while RBDs from civet and sporadic 2004 human isolates (GD03T0013, GZ0402) show weaker binding, consistent with their secondary civet origin and limited transmission^31,32^. SARS-CoV-1-related bat virus RBDs bind to human ACE2, in many cases with higher affinity than SARS-CoV-1 itself.

Binding to civet ACE2 was only detected within the SARS-CoV-1 clade whereas pangolin ACE2 binding is strongest within the SARS-CoV-2 clade, consistent with viruses isolated from civet or pangolin partitioning specifically within each of these clades. Mice are not a natural host of sarbecoviruses, and RBDs from the SARS-CoV-1 and SARS-CoV-2 clades only bind mouse ACE2 sporadically, typically with modest to weak affinity. The highest binding affinity for mouse ACE2 is found in the cluster of RBDs related to RsSHC014, which can mediate infection and pathogenesis in mice^19^.

Binding to ACE2 from *R. affinis* and particularly *R. sinicus* bats varies sharply among strains in the SARS-CoV-1 and SARS-CoV-2 clades, consistent with an evolutionary arms race driving ACE2 variation in *Rhinolophus* bats^28,29^. The two *R. sinicus* bat ACE2 alleles that we tested only interact with SARS-CoV-1 isolates and the bat RsSHC014-cluster RBDs. No binding to these *R. sinicus* ACE2s was detected in the SARS-CoV-2 clade, consistent with the fact that no viruses in this clade have been isolated from *R. sinicus*. In contrast, we detected strong binding to both *R. affinis* ACE2 alleles among many RBDs in the SARS-CoV-1 and SARS-CoV-2 clades. However, the RBDs of the two viruses in our panel sampled from *R. affinis* bound only modestly (LYRa11) or very weakly (RaTG13) to the *R. affinis* alleles that we tested.

Strikingly, we detected binding to *R. affinis* ACE2s by the RBD of the BtKY72 virus from Kenya^15^ (**Fig. 1b,d**), the first described binding to any ACE2 ortholog for a sarbecovirus outside of Asia^7,14^. To validate this finding, we purified the BtKY72 RBD and *R. affinis* ACE2-Fc fusion proteins recombinantly expressed in human cells, and characterized their interaction using biolayer interferometry (BLI). In agreement with the yeast-display results, the BtKY72 RBD bound strongly to the *R. affinis* 9479 ACE2 and more weakly to the *R. affinis* 787 allele (**Fig. 1e**). Furthermore, 293T cells transfected with the *R. affinis* 9479 (but not the 787) ACE2 allele supported entry of vesicular stomatitis virus (VSV) particles pseudotyped with the full-length BtKY72 spike, thereby demonstrating ACE2 is a bona fide entry receptor for this virus (**Fig. 1f** and **Extended Data Fig. 3**). The geographic range of *R. affinis* does not extend outside of Asia^17^, but this result suggests that BtKY72 may bind ACE2 orthologs of bats found in Africa, though the full range of non-Asian bat species that harbor sarbecoviruses and their ACE2 sequences are underexplored^14–16,33^.

We did not detect ACE2 binding by any of the “Clade 2” RBDs. Nine of the 23 Clade 2 RBDs in our tree were sampled from *R. sinicus*, including in the same caves—and even co-infecting the same *R. sinicus* bats^8^—as ACE2-utilizing SARS-CoV-1-related RBDs. Sarbecoviruses with Clade 2 RBDs have also been found across a more diverse array of bat species harboring ACE2s that were not evaluated here, including *R. ferrumequinum, R. macrotis, R. malayanus, R. pusillus, R. pearsonii*, and occasional non-rhinolophid bats^8,9,17,34^. Prior experiments with Clade 2 RBDs have demonstrated a lack of binding to *R. pearsonii*^21^ and human^7,8,12,21^ ACE2, which together with two large deletions in Clade 2 RBDs at the ACE2 interface^7,8,14^, has led to the hypothesis that this clade utilizes some unidentified alternative receptor. Our results are consistent with this hypothesis, though we cannot rule out that these RBDs bind other ACE2 orthologs that have not yet been tested.

### Ancestral origins of sarbecovirus ACE2 binding

Our finding that the BtKY72 RBD binds ACE2 suggests that ACE2 binding was present in the ancestor of all sarbecoviruses prior to the split of Asian and non-Asian RBD clades (**Fig. 2a**). To test this hypothesis, we used ancestral sequence reconstruction^35^ to infer plausible sequences representing ancestral nodes on the sarbecovirus RBD phylogeny (**Fig. 2a** and **Extended Data Fig. 4a**). We evaluated ACE2 binding in the most probable reconstructed ancestral sequences (**Fig. 2b** and **Extended Data Fig. 4b**) and in alternative reconstructions that incorporate statistical or phylogenetic ambiguities inherent to ancestral reconstruction (**Extended Data Fig. 5**). Consistent with the distribution of ACE2 binding among extant sarbecoviruses, the reconstructed ancestor of all sarbecovirus RBDs (AncSarbecovirus) bound the *R. affinis* 9479 ACE2 allele (**Fig. 2b)**. Broader ACE2 binding (including to human ACE2) was acquired on the branch from AncSarbecovirus to the ancestor of the three Asian sarbecoviruses RBD clades (AncAsia). ACE2 binding was then lost along the branch to the Clade 2 ancestor, due to the combination of 48 amino-acid substitutions and 2 deletions at the ACE2-interface that occurred along this branch (**Fig. 2c**).

**Figure 2.**
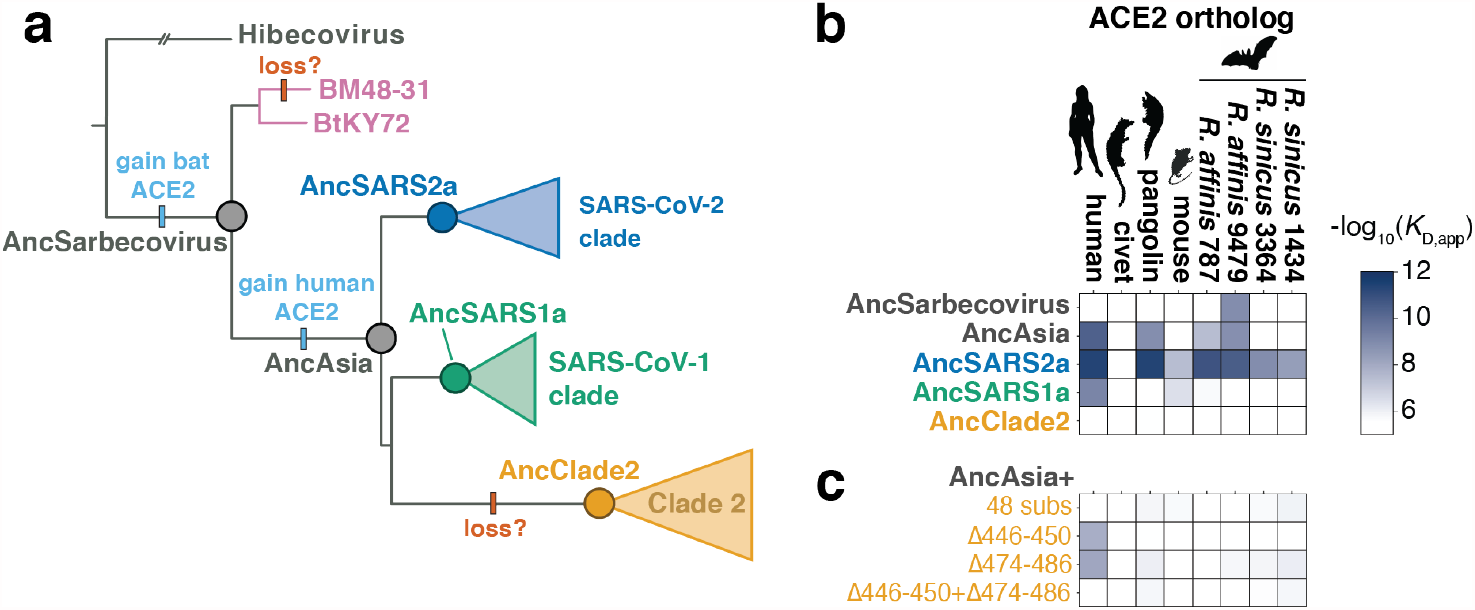
Ancestral origins of sarbecovirus ACE2 binding. **a**, Clade-collapsed RBD phylogeny. Circles represent nodes at which ancestral sequences were inferred. Bars indicate putative gains and losses in ACE2 binding. **b**, ACE2 binding affinities of ancestrally reconstructed RBDs. Additional details in **Extended Data Figs. 4, 5. c**, ACE2 binding affinities of AncAsia plus introduction of the 48 substitutions or two sequence deletions that occurred on the phylogenetic branch leading to AncClade2.

This evolutionary history of ACE2 binding is robust to some but not all explorations of uncertainty in our phylogenetic reconstructions^36,37^. The key phenotypes represented in **Fig. 2b** are robust to uncertainties in the topology of the RBD phylogeny (**Extended Data Fig. 5a,b**) or possible recombination within the RBD impacting the cluster of RBDs related to RsSHC014 (**Extended Data Fig. 5c-f**). However, statistical uncertainty in the reconstructed identity of some ACE2-contact positions impacts our inferences, with some reasonably plausible “second-best’ reconstructed states altering ancestral phenotypes (**Extended Data Fig. 5b**). Nonetheless, our hypothesis of an ancestral origin of sarbecovirus ACE2 binding is supported by the most plausible ancestral reconstructions as well as the distribution of ACE2 binding among directly sampled sarbecovirus RBDs in **Fig. 1b**.

### Evolvability of novel ACE2 binding capabilities

To explore how easily RBDs can acquire ACE2 binding via single amino-acid mutations, we constructed mutant libraries in 14 RBD backgrounds spanning the RBD phylogeny. In each background, we introduced all single amino-acid mutations at six key ACE2-contact positions (SARS-CoV-2 residues L455, F486, Q493, S494, Q498, and N501, **Fig. 3a**; we use SARS-CoV-2 numbering for mutations in all homologs below), encompassing sites of key adaptations or affinity-enhancing mutations described in prior work on SARS-CoV-2 and SARS-CoV-1^12,38^. We recovered nearly all the intended 1,596 mutations, and measured binding of each mutant RBD to each ACE2 ortholog via high-throughput titrations as described above (**Extended Data Fig. 6**).

**Figure 3.**
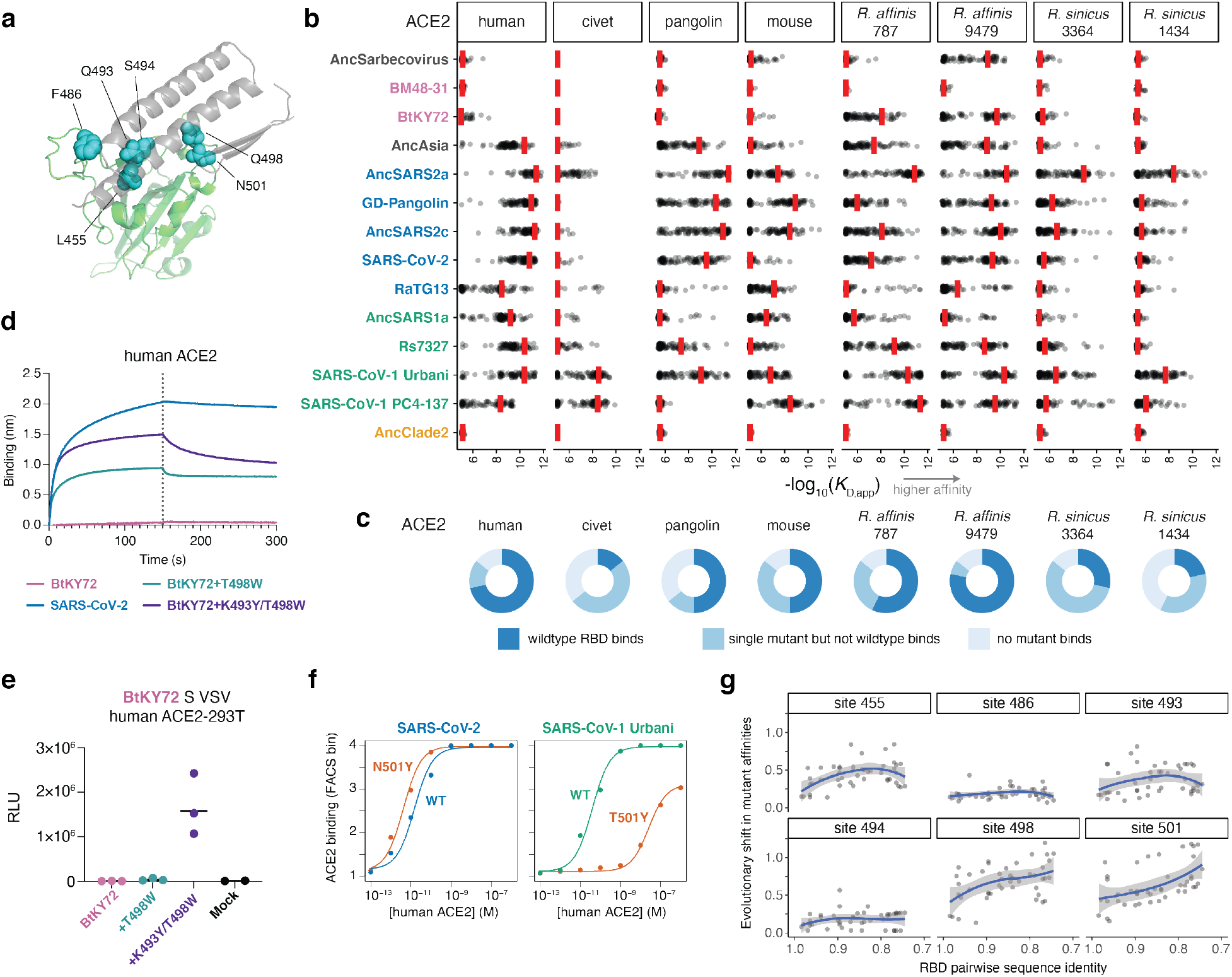
Evolutionary plasticity of ACE2 binding. **a**, Structural context of positions targeted for mutagenesis. RBD as green cartoon, with blue spheres indicating the SARS-CoV-2 residues targeted via mutagenesis. The RBD-interacting region of ACE2 is shown in gray. **b**, Mutational scanning measurements. For each facet, red bars mark the binding affinity of the parental RBD for indicated ACE2. Each point is the affinity of one of the mutations at the six targeted positions. See **Extended Data Fig. 6** for mutation-level measurements. **c**, Pie charts indicating the fraction of the 14 RBD backgrounds for which the parental RBD binds the indicated ACE2 ortholog with -log_10_(*K*_D,app_) > 7, a single mutant binds but the wildtype does not, or no tested mutants bind. **d**, Binding of 1 µM human ACE2-Fc to the biotinylated parental SARS-CoV-2 RBD, BtKY72 RBD or mutant BtKY72 RBDs immobilized at the surface of streptavidin biosensors. Data representative of three assays using independent preparations of RBD (biological triplicate). **e**, Entry of BtKY72 spike-pseudotyped VSV (parental or mutant) in 293T cells stably expressing human ACE2. Each point represents the mean of technical triplicates in assays performed with independent preparation of pseudoviral particles (biological replicate). Geometric mean is indicated by horizontal line. Mock, VSV particles produced in cells in which no spike gene was transfected. Western blot of pseudotyped particles in **Extended Data Fig. 3a**, and entry into 293T cells lacking ACE2 in **Extended Data Fig. 3c. f**, Titration curves showing how mutating site 501 (SARS-CoV2-2 numbering) to tyrosine increases the affinity of the SARS-CoV-2 RBD for human ACE2, but decreases affinity in the SARS-CoV-1 Urbani RBD (**Extended Data Fig. 6**). **g**, Epistatic turnover in mutation effects. Each point represents, for a pair of RBDs, the mean absolute error (residual) in their correlated mutant affinities for human ACE2 binding (**Extended Data Fig. 8a**) versus their pairwise amino acid sequence identity. Correlations computed only for RBD pairs where the parental RBDs bind with -log_10_(*K*_D,app_) > 7. Blue line and shaded gray indicates LOESS mean and 95% CI trendline. Plots incorporating affinity measurements across all ACE2 ligands shown in **Extended Data Fig. 8b**.

The results show that ACE2 binding is a remarkably evolvable trait (**Fig. 3b,c** and **Extended Data Fig. 6**). In virtually all cases in which a parental RBD binds a particular ACE2, there are single amino-acid mutations that improve binding by >5-fold. Therefore, ACE2 binding can be easily enhanced via mutation, which may facilitate the frequent host jumps seen among sarbecoviruses^39^. Notably, our data on mouse ACE2 binding could inform the development of mouse-adapted sarbecovirus strains for *in vivo* studies^19,20,40,41^, including potentially safer strains that bind to mouse but not human ACE2 (see **Extended Data Fig. 7** for details).

In the majority of cases where an RBD does not bind a particular ACE2 ortholog, single mutations can confer low to moderate binding affinity (**Fig. 3b,c**). The only exceptions are BM48-31 and AncClade2, for which none of the tested mutations enabled binding to any of the ACE2s. We found that the mutation K493Y in AncSarbecovirus enables binding to human ACE2 (**Fig. 3b** and **Extended Data Fig. 6**), although this particular mutation did not occur on the branch to AncAsia where we inferred human ACE2 binding was historically acquired, illustrating the existence of multiple evolutionary paths to human ACE2 binding. We identified single mutations at positions 493, 498, and 501 that enable the BtKY72 RBD to bind human ACE2 (**Fig. 3b** and **Extended Data Fig. 6**), suggesting human ACE2 binding is evolutionarily accessible in this lineage.

We validated that mutations K493Y and T498W enable the RBD of the African sarbecovirus BtKY72 to interact with human ACE2 using purified recombinant proteins. Binding to human ACE2-Fc is not detectable with the parental BtKY72 RBD using BLI, but is conferred by T498W and enhanced for the K493Y/T498W double mutant (**Fig. 3d**). To evaluate if the observed binding translated into cell entry, we generated VSV particles pseudotyped with the wildtype or mutant BtKY72 spikes and tested entry in 293T cells expressing human ACE2^42^. We detected robust spike-mediated entry for the K493Y/T498W double mutant but not the T498W single mutant (**Fig. 3e** and **Extended Data Fig. 3**), confirming the evolvability of human ACE2 binding in this African sarbecovirus lineage.

Last, we explored how the mutations that enhance ACE2 binding differ among sarbecovirus backgrounds, reflecting epistatic turnover in mutation effects^12,43^. For example, the N501Y mutation increases human ACE2 binding affinity for SARS-CoV-2 where it has risen in frequency among variants of concern^44^, but the homologous mutation in the SARS-CoV-1 RBD (position 487) is highly deleterious for human ACE2 binding (**Fig. 3f**). More broadly, variation in mutant effects increases as RBD sequences diverge (**Fig. 3g** and **Extended Data Fig. 8**). However, the rate of this epistatic turnover varies across positions—for example, the effects on human ACE2 binding for mutations at positions 486 and 494 remain relatively constant across sequence backgrounds, while variability in effects of mutations at positions 498 and 501 increases substantially as RBDs diverge (**Fig. 3g**).

### Unsampled sarbecovirus lineages are likely ACE2-utilizing

An important consequence of our conclusion that ACE2 binding is an ancestral sarbecovirus trait with plastic evolutionary potential is that unsampled sarbecoviruses lineages are likely capable of binding ACE2 and evolving to bind human ACE2, unless these traits have been specifically lost as occurred in Clade 2. To test this idea, we investigated newly described sarbecoviruses reported after the initiation of our study (**Fig. 4a**). This includes new sarbecoviruses from Africa^14^, as well as a new RBD lineage represented by RsYN04 from a *R. stheno* bat in Yunnan, China^17^, which branches separately from the four RBD clades previously described.

**Figure 4.**
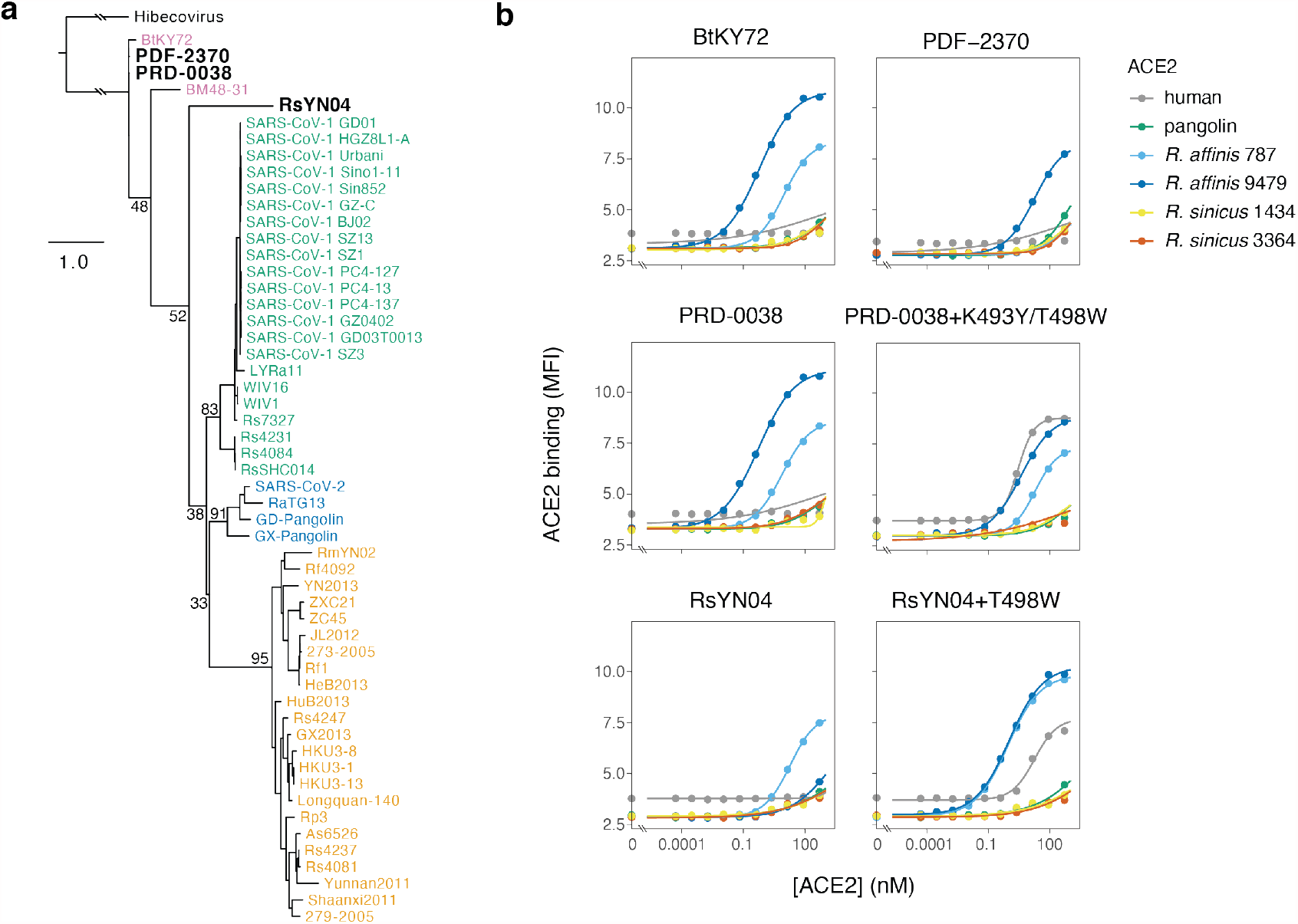
Newly sampled sarbecovirus lineages bind ACE2. **a**, Phylogenetic placement of newly described sarbecovirus RBDs. New sequences in black bold letters, other sequences colored as in **Fig. 1a. b**, Binding curves for new RBDs and candidate mutations that confer human ACE2 binding. Measurements performed with yeast-displayed RBDs and purified dimeric ACE2 proteins, measured by flow cytometry. Data from a single experimental replicate.

We determined the affinity of these RBDs for a panel of ACE2 orthologs using our yeast-display platform (**Fig. 4b**). Like BtKY72, RBDs from the African sarbecoviruses PDF-2380 and PRD-0038^14^ bind *R. affinis* ACE2s, and the K493Y/T498W double mutant confers human ACE2 receptor binding on the PRD-0038 RBD (**Fig. 4b**) as it does for BtKY72 (**Fig. 3e**). The RsYN04 RBD binds to *R. affinis* 787 ACE2, as was recently shown for the closely related RaTG15^45^. The RsYN04 RBD can also acquire binding to human ACE2 through the single T498W mutation. These results illustrate that the ancestral traits of ACE2 binding and ability to evolve human ACE2 binding are maintained in geographically and phylogenetically diverse sarbecovirus lineages, including ones just beginning to be described^14,15,17,45^.

## Discussion

Our experiments reveal that binding to bat ACE2 is an ancestral trait of sarbecoviruses that is also present in viruses from outside of Asia^14,15^. Binding to human ACE2 arose in the common ancestor of SARS-CoV-1- and SARS-CoV-2-related RBDs prior to their divergence. Binding to the ACE2 orthologs we tested was then lost on the branch leading to the Clade 2 RBDs, which must bind an alternative receptor or ACE2 orthologs not tested in our work. These results expand our understanding of the phylogenetic breadth of ACE2 binding and imply that unsampled RBD lineages in the phylogenetic interval between BtKY72 and SARS-CoV-1 or SARS-CoV-2 are likely ACE2 utilizing.

Our work also shows that ACE2 binding is a highly evolvable trait of sarbecovirus RBDs. For every ACE2-binding RBD we studied, there were single amino-acid mutations that enhanced affinity for ACE2 orthologs the RBD could already bind or conferred binding to new ACE2 orthologs from different species. Sarbecoviruses are infamous because two different strains have caused human outbreaks, and host jumps are probably far more common among the wide diversity of bats that are naturally infected with these viruses^8,17,39^. The evolutionary plasticity of RBD ACE2 binding is therefore likely a key contributor to the ecological dynamics of sarbecoviruses. However, because the effects of RBD mutations on ACE2 binding can differ across sarbecovirus backgrounds, it is not trivial to predict an RBD’s ACE2 binding properties from sequence alone. Therefore, the high-throughput approach we have used here, which enables rapid and comprehensive measurement of ACE2 binding affinities without requiring work with live viruses, can aid efforts to understand the evolutionary diversity and dynamics of sarbecoviruses.

## Acknowledgements

We thank the Fred Hutch Flow Cytometry and Genomics facilities and the Scientific Computing group supported by ORIP grant S10OD028685. We thank Dr. Shi Zhengli (Wuhan Institute of Virology) for sharing *R. sinicus* and *R. affinis* ACE2 sequences from a preprint prior to their formal publication. We thank Hideki Tani (University of Toyama) for providing the reagents necessary for preparing VSV pseudotyped viruses. We thank Stephen Goldstein and Alina Chan for helpful discussions. This study was supported by the National Institute of Allergy and Infectious Diseases (DP1AI158186 and HHSN272201700059C to DV, and R01AI141707 to JDB), the National Institute of General Medical Sciences (R01GM120553 to DV and 5T32GM008268-32 to SKZ) a Pew Biomedical Scholars Award (DV), Investigators in the Pathogenesis of Infectious Disease Awards from the Burroughs Wellcome Fund (DV and JDB), Fast Grants (DV), and the Bill & Melinda Gates Foundation (OPP1156262 to DV and INV-004949 to JDB). TNS is an HHMI Fellow of the Damon Runyon Cancer Research Foundation. JDB is an Investigator of the Howard Hughes Medical Institute.

## Author contributions

Conceptualization: TNS, SKZ, DV, JDB. Phylogenetics: TNS. Yeast-display methodology: TNS, AJG. Yeast-display experiments: TNS. BLI measurements: SKZ. Pseudovirus entry assays: SKZ, ACW. Data analysis: TNS. Supervision: DV, JDB. Writing – original draft: TNS, SKZ, DV, JDB. Writing – review and editing: all authors

## Competing interests

JDB consults for Moderna on viral evolution and epidemiology and Flagship Labs 77 on deep mutational scanning. JDB may receive a share of IP revenue as an inventor on a Fred Hutchinson Cancer Research Center-optioned technology/patent (application WO2020006494) related to deep mutational scanning of viral proteins. DV is a consultant for Vir Biotechnology Inc. The Veesler laboratory has received an unrelated sponsored research agreement from Vir Biotechnology Inc.

## Additional Information

Correspondence and requests for materials should be addressed to Tyler N Starr, David Veesler, and Jesse D Bloom

## METHODS

### Phylogenetics and ancestral sequence reconstruction

All steps of bioinformatic analysis, including specific programmatic commands, alignments, raw data, and output files can be found on GitHub: https://github.com/jbloomlab/SARSr-CoV_homolog_survey/tree/master/RBD_ASR.

A panel of unique sarbecovirus RBD sequences was assembled incorporating the RBD sequences curated by Letko et al.^7^, all unique RBD sequences among SARS-CoV-1 human and civet strains reported by Song et al.^32^, and recently reported sarbecoviruses BtKY72^15^, RaTG13^2^, GD-Pangolin-CoV (consensus RBD sequence reported in Fig. 3a of Lam et al.^25^) and GX-Pangolin-CoV^25^ (P2V, ambiguous nucleotide in codon 515 (SARS-CoV-2 numbering) was resolved to retain amino acid F515, which is conserved across all other sarbecoviruses). We also incorporated newly described sarbecovirus sequences RsYN04^17^, PDF-2370 and PRD-0038^14^ into updated phylogenies and functional work after the initiation of our study. The Hibecovirus sequence Hp-BetaCoV/Zhejiang2013 (Genbank: KF636752) was used to root the sarbecovirus phylogeny. For **Extended Data Fig. 1**, additional betacoronavirus outgroups were included in rooting. All virus names and sequence accessions or citations are provided on GitHub: https://github.com/jbloomlab/SARSr-CoV_homolog_survey/blob/master/RBD_ASR/RBD_accessions.csv. We thank all sequence contributors, including contributors to GISAID: https://github.com/jbloomlab/SARSr-CoV_homolog_survey/tree/master/RBD_ASR/gisaid.

Amino acid sequences were aligned by mafft (version 7.471)^46^ with a gap opening penalty of 4.5. RBD sequences were subsetted from spike alignments according to our domain boundary defined for SARS-CoV-2 (Wuhan-Hu-1 Genbank: MN908947, residues N331-T531). Nucleotide alignments were constructed from amino acid alignments using PAL2NAL (version 14)^47^. Phylogenies were inferred with RAxML (version 8.2.12)^48^ using the LG+Γ substitution model for amino acid sequence alignments or GTR+Γ with separate data partitions applied to the first, second, and third codon positions for nucleotide sequence alignments. Constraint files specifying specific clade relationships (but free topologies within clades) were used to test specific alternate topologies in **Extended Data Fig. 5a**.

Marginal likelihood ancestral sequence reconstruction was performed with FastML (version 3.11)^49^ using the amino acid sequence alignment, the maximum likelihood nucleotide tree topology from RAxML, the LG+Γ substitution matrix, re-optimization of branch lengths, and FastML’s likelihood-based indel reconstruction model. The maximum *a posteriori* (MAP) ancestral sequences at nodes of interest were determined from the marginal reconstructions as the string of amino acids at each alignment site with the highest posterior probability, censored by deletions as inferred from the indel reconstruction. To test the robustness of ancestral phenotypes to statistical uncertainty in reconstructed ancestral states, we also constructed “alt” ancestors in which all second-most-probable states with posterior probability > 0.2 were introduced simultaneously^36^.

To identify potential recombination breakpoints within the RBD alignment, we used GARD (version 0.2)^50^, which identified a possible recombination breakpoint (**Extended Data Fig. 5c**) that produces two alignment segments exhibiting phylogenetic incongruence with a gain in overall likelihood sufficient to justify the duplication of phylogenetic parameters (ΔAIC -85). To determine the impact of this possible recombination on ancestral sequence reconstructions, the alignment was split into separate segments at the proposed breakpoint. Phylogenies were inferred and ancestral sequences reconstructed on separate segments as described above, and reconstructed ancestral sequences at matched nodes for each segment were concatenated, as shown in **Extended Data Fig. 5e**.

### RBD library construction

Genes encoding all 73 unique extant and ancestral RBD amino acid sequences were ordered from Twist Bioscience, Genscript, and IDT. Gene sequences are provided on GitHub: https://github.com/jbloomlab/SARSr-CoV_homolog_survey/blob/master/RBD_ASR/parsed_sequences/RBD_sequence_set_annotated.csv. Genes were cloned in bulk into the pETcon yeast surface-display vector (plasmid 2649) as described by Starr et al.^12^. As described in this prior publication, randomized N16 barcodes were appended via PCR downstream from RBD coding sequences. RBD sequences were pooled and barcoded in two independently processed replicates. The pooled, barcoded parental RBD libraries were electroporated into *E. coli* and plated at an estimated bottleneck of ∼22,000 cfu, yielding an estimated ∼300 barcodes per parental RBD within each library replicate.

In parallel, we cloned site saturation mutagenesis libraries of six positions in select RBD backgrounds. The positions targeted correspond to SARS-CoV-2 positions 455, 486, 493, 494, 498, and 501. The RBD-indexed position targeted in each background is provided on GitHub: https://github.com/jbloomlab/SARSr-CoV_homolog_survey/blob/master/RBD_ASR/parsed_sequences/RBD_sequence_set_annotated.csv. Precise site saturation mutagenesis pools were produced by Genscript, provided as plasmid libraries. Failed positions in the Genscript mutagenesis libraries (all six positions in SARS-CoV-1 Urbani, position 494 in SARS-CoV-2, and position 455 in RaTG13 and GD-Pangolin) or backgrounds chosen for mutagenesis subsequent to initial library design (BtKY72) were produced in-house via PCR-based mutagenesis using NNS degenerate mutagenic primers followed by Gibson Assembly of mutagenized fragments. In duplicate, mutant libraries were pooled and N16 barcodes were appended downstream from the RBD coding sequence. The pooled, barcoded mutant libraries were electroporated into *E. coli* and plated at a target bottleneck corresponding to an average of 20 barcodes per mutant within each library replicate.

Colonies from bottlenecked transformation plates were scraped and plasmid purified. Parental RBD and mutant pools were combined at ratios corresponding to expected barcode diversity, yielding the two separately barcoded library replicates used in high-throughput experiments. Plasmid libraries were transformed into yeast (AWY101 strain^51^) according to the protocol of Gietz and Schiestl^52^, transforming 10 µg of plasmid at 10× scale.

### PacBio sequencing and analysis

As described by Starr et al.^12^, PacBio sequencing was used to acquire long sequence reads spanning the N16 barcode and RBD coding sequence. PacBio sequencing constructs were prepared from library plasmid pools via NotI digestion and gel purification, followed by SMRTbell ligation. Each library was sequenced across three SMRT Cells on a PacBio Sequel using 20-hour movie collection times. PacBio circular consensus sequences (CCSs) were generated from subreads using the ccs program (version 5.0.0), requiring 99.9% accuracy and a minimum of 3 passes. The resulting CCSs are available on the NCBI Sequence Read Archive, BioSample SAMN18316101.

CCSs were processed using alignparse (version 0.1.6)^53^ to identify the RBD target sequence, call any mutations, and determine the associated N16 barcode sequence, requiring no more than 18 nucleotide mutations from the intended target sequence, an expected 16-nt length barcode sequence, and no more than 3 mismatches across the sequenced portions of the vector backbone.

We next used processed CCSs to link each barcode to the associated RBD sequence. We first filtered sequences with ccs-determined accuracies of <99.99% or indels. The empirical sequencing accuracy estimated by comparing RBD variants associated with barcode sequences sampled across multiple CCSs (https://jbloomlab.github.io/alignparse/alignparse.consensus.html#alignparse.consensus.empirical_accuracy) was 99.0% and 98.4% in libraries 1 and 2, respectively. For barcodes sampled across multiple CCSs, we derived consensus RBD variant sequences, discarding barcodes where CCSs with identical barcodes exhibited >1 point mutation or >2 indels, or where >10% or >25% of CCSs with an identical barcode contain a secondary non-consensus mutation or indel, respectively. The CCS processing pipeline is available on GitHub: https://github.com/jbloomlab/SARSr-CoV_homolog_survey/blob/master/results/summary/process_ccs.md. The final barcode-variant lookup table, which links each N16 barcode with its associated RBD sequence, is available on GitHub: https://github.com/jbloomlab/SARSr-CoV_homolog_survey/blob/master/results/variants/nucleotide_variant_table.csv.

### ACE2 proteins for yeast display assays

Recombinant dimeric ACE2 proteins were purchased or produced from commercial sources. Recombinant human ACE2 (Uniprot: Q9BYF1-1) was purchased from ACROBiosystems (AC2-H82E6), consisting of residues 18-740 spanning an intrinsic dimerization domain, followed by a His tag and biotinylated Avitag used for downstream detection. Civet (*Paguma larvata*) ACE2 (Uniprot: Q56NL1-1) was purchased from ACROBiosystems (AC2-P5248), consisting of residues 18-740 spanning an intrinsic dimerization domain, with an N-terminal His tag used for downstream detection. Mouse (*Mus musculus*) ACE2 (Uniprot: Q8R0I0-1) was purchased from Sino Biological (50249-M03H), consisting of residues 18-740 spanning an intrinsic dimerization domain, followed by a His tag and human IgG1 Fc domain used for downstream detection.

The remaining ACE2s were produced by Genscript. Specifically, pangolin (*Manis javanica*, Genbank: XP_017505746.1), *R. affinis* 787 (Genbank: QMQ39222), *R. affinis* 9479 (Genbank: QMQ39227), *R. sinicus* 3364 (Genbank: QMQ39219), and *R. sinicus* 1434 (Genbank: QMQ39216) ACE2 residues 19-615 were cloned with a C-terminal human IgG1 Fc domain for dimerization and downstream detection. pcDNA3.4 expression plasmids were transfected into HD 293F cells for protein expression. ACE2-Fc fusions were purified from day six culture supernatants via Fc-tag affinity purification.

### Library measurements of RBD expression and RBD+ enrichment

Transformed yeast library aliquots were grown overnight in a shaker at 30°C in SD-CAA media (6.7 g/L Yeast Nitrogen Base, 5.0 g/L Casamino acids, 1.065 g/L MES, and 2% w/v dextrose, pH 5.3). To induce RBD expression, yeast were washed and resuspended in SG-CAA+0.1%D media (6.7 g/L Yeast Nitrogen Base, 5.0 g/L Casamino acids, 1.065 g/L MES, 2% w/v galactose, and 0.1% w/v dextrose, pH 5.3) at initial OD600 0.67, and incubated at room temperature for 16-18 hr with mild agitation.

For each library, 45 OD of induced culture was washed twice with PBS-BSA (0.2 mg/mL), and RBD surface expression was labeled via a C-terminal c-Myc tag with 1:100 diluted FITC-conjugated chicken anti-c-Myc antibody (Immunology Consultants Lab, CMYC-45F) in 3mL PBS-BSA. Labeled cells were washed twice in PBS-BSA, and resuspended in PBS for FACS.

Yeast library sorting experiments were conducted on a BD FACSAria II with FACSDiva software (version 8.0.2). For high-throughput measurements of RBD expression levels, cells were gated for single cells (**Extended Data Fig. 2b**), and partitioned into four bins of FITC fluorescence (**Extended Data Fig. 2c**), where bin 1 captures 99% of unstained cells, and bins 2-4 split the remaining library population into tertiles. Cells were sorted into 5mL tubes pre-wet with 1mL of SD-CAA with 1% BSA. We recovered ∼8 million cells per library across the four bins. Sorted cells were resuspended to 2e6 cells/mL in fresh SD-CAA with 1:100 penicillin-streptomycin, and grown overnight at 30°C. Plasmid was purified from post-sort yeast samples of <4e7 cells per miniprep column using the Zymo Yeast Miniprep II kit (D2004) according to manufacturer instructions, with the addition of an extended (>2 hr) Zymolyase treatment and a -80°C freeze/thaw cycle prior to cell lysis. N16 barcodes were PCR amplified from each plasmid aliquot as described in Starr et al.^12^ and submitted for Illumina HiSeq 50bp single end sequencing.

To enrich properly expressing RBD variants for downstream titration experiments, we also sorted ∼2e7 cells per library using the RBD+ (FITC+) bin shown in **Extended Data Fig. 2b**). RBD+-enriched populations were resuspended to 1e6 cells/mL for overnight outgrowth, and frozen -80°C in 9 OD aliquots for subsequent titration experiments.

A pool of mutants that were added after the first set of experiments (mutations at position 455 in RaTG13 and GD-Pangolin, and mutations at all six positions in BtKY72) were not RBD+ enriched and were not part of the bulk expression Sort-seq measurement, but were pooled with the RBD+-enriched population of the primary libraries for subsequent titration assays.

### Library measurements of ACE2 binding affinities

For high-throughput measurements of ACE2 binding affinities, yeast libraries were induced for RBD expression as described above. Induced cultures were aliquoted at 8 OD per titration sample and washed twice with PBS-BSA. Cells were resuspended across a range of ACE2 concentrations from 1e-6 to 1e-13 M in 1 M intervals, plus a 0 M ACE2 concentration. Samples were incubated overnight at room temperature with mild agitation. Samples were washed twice in ice-cold PBS-BSA, and resuspended in 1mL secondary label (1:100 Myc-FITC, and 1:200 PE-conjugated streptavidin (Thermo Fisher S866) for human ACE2, 1:200 iFluor647-conjugated mouse anti-His (Genscript A01802) for civet ACE2, and 1:200 PE-conjugated goat anti-human IgG (Jackson ImmunoResearch Labs 109-115-098) for all other Fc-tagged ACE2 ligands), and incubated for 1 hour on ice. Cells were washed twice with PBS-BSA and resuspended in PBS for FACS.

Titration samples were binned for single, RBD-expressing cells (**Extended Data Fig. 2b**), which were then partitioned into four bins on the basis of ACE2 binding (**Extended Data Fig. 2d**). At each concentration, a minimum of 5e6 cells were collected across the four bins. Sorted cells were resuspended in 1mL SD-CAA with 1:100 penicillin-streptomycin, and grown overnight at 30°C in deep well plates. Plasmid aliquots from each population were purified with the Zymo Yeast 96-Well Miniprep kit (D2005) according to manufacturer instructions, with the addition of an extended (>2 hr) Zymolyase treatment and a -80°C freeze/thaw cycle prior to cell lysis. N16 barcodes were PCR amplified from each plasmid aliquot as described in Starr et al.^12^ and submitted for Illumina HiSeq 50bp single end sequencing.

For the pool of mutants that were added after the first set of experiments (mutations at position 455 in RaTG13 and GD-Pangolin, and mutations at all six positions in BtKY72), duplicate titrations were already conducted with the primary pool for human ACE2 and *R. affinis* 787 ACE2. Titrations with this smaller library sub-pool with these ACE2 ligands were conducted as above, but scaled to 1.6 OD per sample, collecting >1 million cells per concentration.

### Illumina barcode sequencing analysis

Demultiplexed sequence reads (available on the NCBI Sequence Read Archive, BioSample SAMN20174027) were aligned to library barcodes as determined from PacBio sequencing using dms_variants (version 0.8.5), yielding a count of the number of times each barcode was sequenced within each FACS bin. Read counts within each FACS bin were downweighted by the ratio of total reads from a bin compared to the number of cells that were actually sorted into that bin. The table giving downweighted counts of each barcode in each FACS bin is available on GitHub: https://github.com/jbloomlab/SARSr-CoV_homolog_survey/blob/master/results/counts/variant_counts.csv.

We estimated the RBD expression level of each barcoded variant based on its distribution of counts across FACS bins and the known log-transformed fluorescence boundaries of each sort bin using a maximum likelihood approach^12,54^, implemented with the fitdistrplus package (version 1.0.14)^55^ in R. Expression measurements were retained for barcodes for which greater than 20 counts were observed across the four FACS bins. The full pipeline for computing per-barcode expression values is described on GitHub: https://github.com/jbloomlab/SARSr-CoV_homolog_survey/blob/master/results/summary/compute_expression_meanF.md.

We estimated the level of ACE2 binding of each barcoded variant at each titration concentration based on its distribution of counts across FACS bins calculated as a simple mean^54^, as described in Starr et al.^12^. We determined the apparent binding constant *K*_D,app_ describing the affinity of each barcoded variant for each ACE2 along with free parameters *a* (titration response range) and *b* (titration curve baseline) via nonlinear least-squares regression using the standard non-cooperative Hill equation relating the mean bin response variable to the ACE2 labeling concentration:

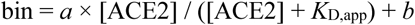

The measured mean bin value at a given ACE2 concentration was excluded from a variant’s curve fit if fewer than 10 counts were observed across the four FACS bins at that concentration. Individual concentration points were also excluded from the curve fit if they demonstrated evidence of bimodality (>40% of counts of a barcode were found in each of two non-consecutive bins 1+3 or 2+4, or >20% of counts of a barcode were found in each of the boundary bins 1+4). To avoid errant fits, we constrained the fit baseline parameter *b* to be between 1 and 1.5, the response parameter *a* to be between 2 and 3, and the *K*_D,app_ parameter to be between 1e-15 and 1e-5. The fit for a barcoded variant was discarded if the average count across all sample concentrations was below 10, or if >20% of sample concentrations were missing due to counts below 10. We also discarded curve fits where the normalized mean square residual (residuals normalized from 0 to 1 relative to the fit response parameter *a*) is >10× the median normalized mean square residual across all titrations with all ACE2s. *K*_D,app_ binding constants were expressed as -log_10_(*K*_D,app_), where higher values indicate higher affinity binding. The full pipeline for computing per-barcode binding affinities is described on GitHub: https://github.com/jbloomlab/SARSr-CoV_homolog_survey/blob/master/results/summary/compute_binding_Kd.md.

To derive our final measurements we collapsed measurements across internally replicated barcodes representing each RBD genotype. For each RBD genotype, we discarded the top and bottom 5% (expression measurements) or 2.5% (titration affinities) of per-barcode measurements, and computed the mean value across remaining barcodes within each library. The correlations in these barcode-averaged measurements between independently barcoded and assayed library replicates are shown in **Extended Data Fig. 2g**. Final measurements were determined as the mean of the barcode-collapsed mean measurements from each replicate. The total number of barcodes collapsed into these final measurements from both replicates are shown in the histograms in **Extended Data Fig. 2f**. Final measurements for an RBD genotype were discarded if the RBD genotype was not sampled with at least one non-filtered barcode in each replicate, or sampled with at least five non-filtered barcodes in a single replicate. The full pipeline for barcode collapsing is described on GitHub: https://github.com/jbloomlab/SARSr-CoV_homolog_survey/blob/master/results/summary/barcode_to_genotype_phenotypes.md. The final processed measurements of expression and ACE2 binding for parental and mutant RBDs can be found on GitHub: https://github.com/jbloomlab/SARSr-CoV_homolog_survey/blob/master/results/final_variant_scores/wt_variant_scores.csv and https://github.com/jbloomlab/SARSr-CoV_homolog_survey/blob/master/results/final_variant_scores/mut_variant_scores.csv.

### lsogenic ACE2 binding assays

For RBDs assayed subsequent to library experiments (**Fig. 4** and **Extended Data Fig. 5f**), RBDs were cloned as isogenic stocks into the 2649 plasmid, sequence verified, and transformed individually into yeast using the LiAc/ssDNA transformation method^56^. Cultures were induced for RBD expression and labeled across ACE2 concentration series as described above, in V-bottom 96-well plates with 0.067 OD yeast per well. ACE2 labeling of RBD+ cells was measured on a BD LSRFortessa X50 flow cytometer and data was processed via FlowJo (version 10). Binding curves of PE (ACE2) mean fluorescence intensity versus ACE2 labeling concentration was fit as above, with the inclusion of a hill coefficient slope parameter *n*.

### Transient expression of R. affinis ACE2-Fc

The *R. affinis* 787 (GenBank: QMQ39222.1) and *R. affinis* 9479 (GenBank: QMQ39227.1) ACE2 ectodomains constructs were synthesized by GenScript and placed into a pCMV plasmid. The domain boundaries for the ectodomain are residues 19-615. The native signal tag was identified using SignalP-5.0 (residues 1-18) and replaced with a N-terminal mu-phosphatase signal peptide. These constructs were then fused to a sequence encoding thrombin cleavage site and a human Fc fragment at the C-terminal end. All ACE2-Fc constructs were produced in Expi293F cells (Thermo Fisher A14527) in Gibco Expi293 Expression Medium at 37°C in a humidified 8% CO2 incubator rotating at 130 rpm. The cultures were transfected using PEI-25K (Polyscience) with cells grown to a density of 3 million cells per mL and cultivated for 4-5 days. Proteins were purified from clarified supernatants using a 1 mL HiTrap Protein A HP affinity column (Cytiva), concentrated and flash frozen in 1x PBS, pH 7.4 (10 mM Na_2_HPO_4_, 1.8 mM KH_2_PO_4_, 2.7 mM KCl, 137 mM NaCl).

### Transient expression of BtKY72 parental and mutant RBDs

BtKY72 RBD construct (BtKY72 S residues 318-520) was synthesized by GenScript into a CMVR plasmid with a N-terminal mu-phosphatase signal peptide and a C-terminal hexa-histidine tag (-HHHHHHHH) joined by a short linker (-GGSS) to a Avi tag (-GLNDIFEAQKIEWHE). BtKY72 mutant constructs T498W (BtKY72 S residue 487) and K493Y/T498W (BtKY72 S residue 482/487) were subcloned by GenScript from the BtKY72 RBD construct. BtKY72 and BtKY72 mutant RBD constructs were produced in Expi293F cells in Gibco Expi293 Expression Medium at 37°C in a humidified 8% CO2 incubator rotating at 130 rpm. The cultures were transfected using PEI-25K with cells grown to a density of 3 million cells per mL and cultivated for 3-5 days. Proteins were purified from clarified supernatants using a 1mL HisTrap HP affinity column (Cytiva), concentrated, and then biotinylated with a commercial BirA kit (Avidity). Proteins were then purified from the BirA enzyme by affinity purification using a 1 mL HisTrap HP affinity column (Cytiva), concentrated, and flash frozen in 1x PBS, pH 7.4.

### Biolayer interferometry

Assays were performed on an Octet Red (Forte Bio) instrument at 30°C with shaking at 1,000 RPM. Streptavidin biosensors were hydrated in water for 10 min prior to a 60 s incubation in 10x Kinetics Buffer (undiluted). Biotinylated RBDs were loaded at 5-10 μg/mL in 10s Kinetics Buffer for 100-600 s prior to baseline equilibration for 120 s in 10x kinetics buffer. Association of ACE2-Fc (dimeric) was performed at 1 µM in 10x Kinetics Buffer. The data were baseline subtracted. The experiments were done with three separate purification batches of BtKY72 RBDs. All RBDs were immobilized to identical levels, i.e. 1 nm shift. The data were plotted in Graph Prism and a representative plot is shown.

### Generation of VSV pseudovirus

The BtKY72 S construct was synthesized by GenScript and cloned into an HDM plasmid with a C-terminal 3X FLAG tag. The BtKY72 mutant S constructs T498W (BtKY72 S residue 487) and K493Y/T498W (BtKY72 S residue 482/487) were subcloned by GenScript from the BtKY72 S construct. Pseudotyped VSV particles were prepared using HEK293T (293T) (ATCC CRL-11268) cells seeded into 10-cm dishes. 293T cells were transfected using Lipofectamine 2000 (Life Technologies) with a S encoding-plasmid in Opti-MEM transfection medium and incubated for 5 hr at 37°C with 8% CO_2_ supplemented with DMEM containing 10% FBS. One day post-transfection, cells were infected with VSV (G*ΔG-luciferase), and after 2 hr, infected cells were washed 5x with DMEM before adding medium supplemented with anti-VSV G antibody (I1-mouse hybridoma supernatant diluted 1:40, from ATCC CRL-2700). Pseudotyped particles were harvested 18-24 hr post-inoculation, clarified from cellular debris by centrifugation at 3000 g for 10 min, concentrated 100x using a 100 MWCO membrane for 10 min at 3000 rpm, and frozen at -80°C. Mock pseudotyped VSV pseudovirus was generated as above but in the absence of S.

### VSV pseudovirus entry assays

HEK293T (293T) cells (ATCC CRL-11268) and 293T cells with stable transfection of human ACE2^42^ were cultured in 10% FBS, 1% PenStrep DMEM at 37°C in a humidified 8% CO2 incubator. Cells were plated into poly-lysine coated 96-well plates. For *R. affinis* ACE2 entry, transient transfection of *R. affinis* ACE2 in 293T cells was done 24 hours prior to infection using Lipofectamine 2000 (Life Technologies) and an HDM plasmid containing full length *R. affinis* ACE2 (synthesized by GenScript) in OPTIMEM. After 5 hr incubation at 37°C in a humidified 8% CO2 incubator, DMEM with 10% FBS was added and cells were incubated at 37°C in a humidified 8% CO2 incubator for 18 hr.

Immediately prior to infection, 293T cells with stable expression of human ACE2, transient expression of *R. affinis* ACE2 or not transduced to express ACE2 were washed with DMEM 3x, then plated with normalized pseudovirus in DMEM. Infection in DMEM was done with cells between 60-80% confluence (human ACE2 293T) or between 80-90% confluence (*R. affinis* ACE2 293T) for 2.5 hr prior to adding FBS and PenStrep to final concentrations of 10% and 1%, respectively. Following 24 hr of infection, One-Glo-EX (Promega) was added to the cells and incubated in the dark for 5 min before reading on a Synergy H1 Hybrid Multi-Mode plate reader (Biotek). Normalized cell entry levels of pseudovirus generated on different days (biological replicates) were plotted in Graph Prism as individual points, and average cell entry across biological replicates was calculated as the geometric mean.

BtKY72 S parental and mutant pseudoviral particle inputs for the above cell entry assays were normalized by spike incorporation quantified via western blot. Detection of S was done with mouse monoclonal ANTI-FLAG M2 antibody (Sigma F3165) and Alexa Fluor 680 AffiniPure Goat Anti-Mouse IgG, light chain specific (Jackson ImmunoResearch Labs 115-625-174). Detection of the VSV backbone was done with Anti-VSV-M [23H12] Antibody (Kerafast EB0011) and Alexa Fluor 680 AffiniPure Goat Anti-Mouse IgG, light chain specific (Jackson ImmunoResearch Labs 115-625-174). A representative blot is shown in **Extended Data Fig. 3a**.

## Data availability

- PacBio circular consensus sequences are available from the NCBI SRA, BioSample SAMN18316101
- Illumina sequences for barcode counting are available from the NCBI SRA, BioSample SAMN20174027
- Table of measurements of ACE2 binding and expression for all parental RBDs is available on GitHub: https://github.com/jbloomlab/SARSr-CoV_homolog_survey/blob/master/results/final_variant_scores/wt_variant_scores.csv
- Table of measurements of ACE2 binding and expression for all single mutant RBDs is available on GitHub: https://github.com/jbloomlab/SARSr-CoV_homolog_survey/blob/master/results/final_variant_scores/mut_variant_scores.csv

## Code availability

- All code for data analysis is available on GitHub: https://github.com/jbloomlab/SARSr-CoV_homolog_survey
- A summary of the computational pipeline and links to individual notebooks detailing steps of analysis is available on Github: https://github.com/jbloomlab/SARSr-CoV_homolog_survey/blob/master/results/summary/summary.md

**Extended Data Figure 1.**
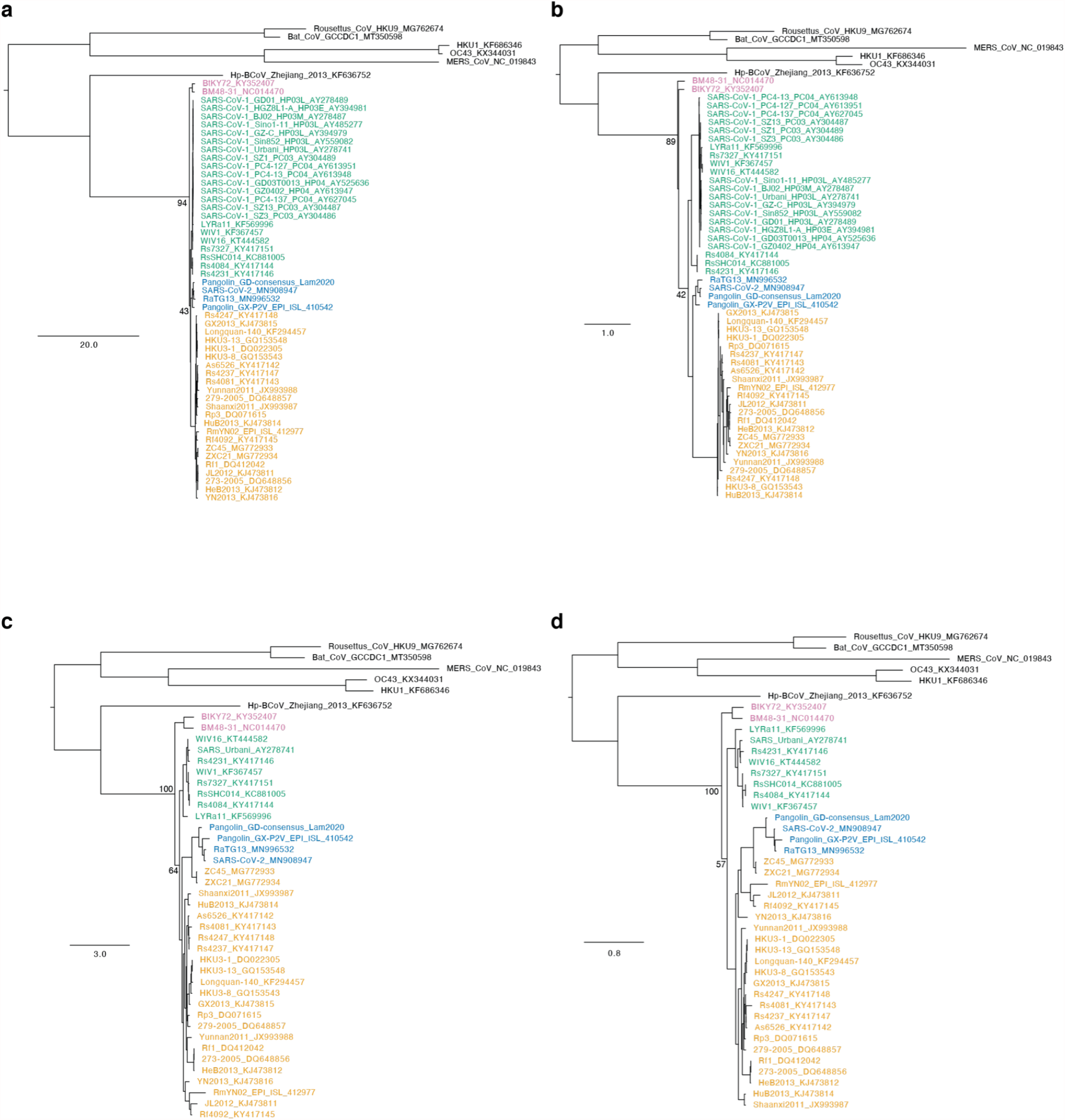
Robustness of the root of the sarbecovirus ingroup. To establish robustness of our conclusion that the first sarbecovirus divergence is between sarbecoviruses from Africa and Europe and those from Asia, we inferred phylogenies based on alignments of RBD (**a,b**) or the full spike gene (**c,d**) and nucleotide (**a, c**) or amino-acid (**b, d**) alignments and substitution models. In all four cases, the first sarbecovirus bipartition is placed between sarbecoviruses in Africa and Europe and those in Asia. The placement of the overall tree root is arbitrary with respect to the relationship among non-sarbecovirus outgroups, but this arbitrary placement does not impact the sarbecovirus ingroup rooting. The primary variations among trees includes a potential paraphyletic separation of BtKY72 and BM48-31 from Europe and Africa such that they do not form a monophyletic clade (**b**; also seen in **Fig. 4a**), and variation in the relationships among the three Asia sarbecovirus clades (whose relationship is also inferred with a very low bootstrap support value in our primary phylogeny in **Fig. 1a**). Known recombination of RBDs with respect to other spike segments among viruses creates incongruencies between spike and RBD trees among Asian sarbecovirus lineages (e.g. ZC45 and ZXC21), though recombination has not been reported among the Africa and Europe spikes and those in Asia. Scale bar, expected number nucleotide or amino-acid substitutions per site. Node labels illustrate bootstrap support values for sarbecovirus and Asia sarbecovirus monophyly. Sequences colored by their RBD clade as in **Fig. 1a**.

**Extended Data Figure 2.**
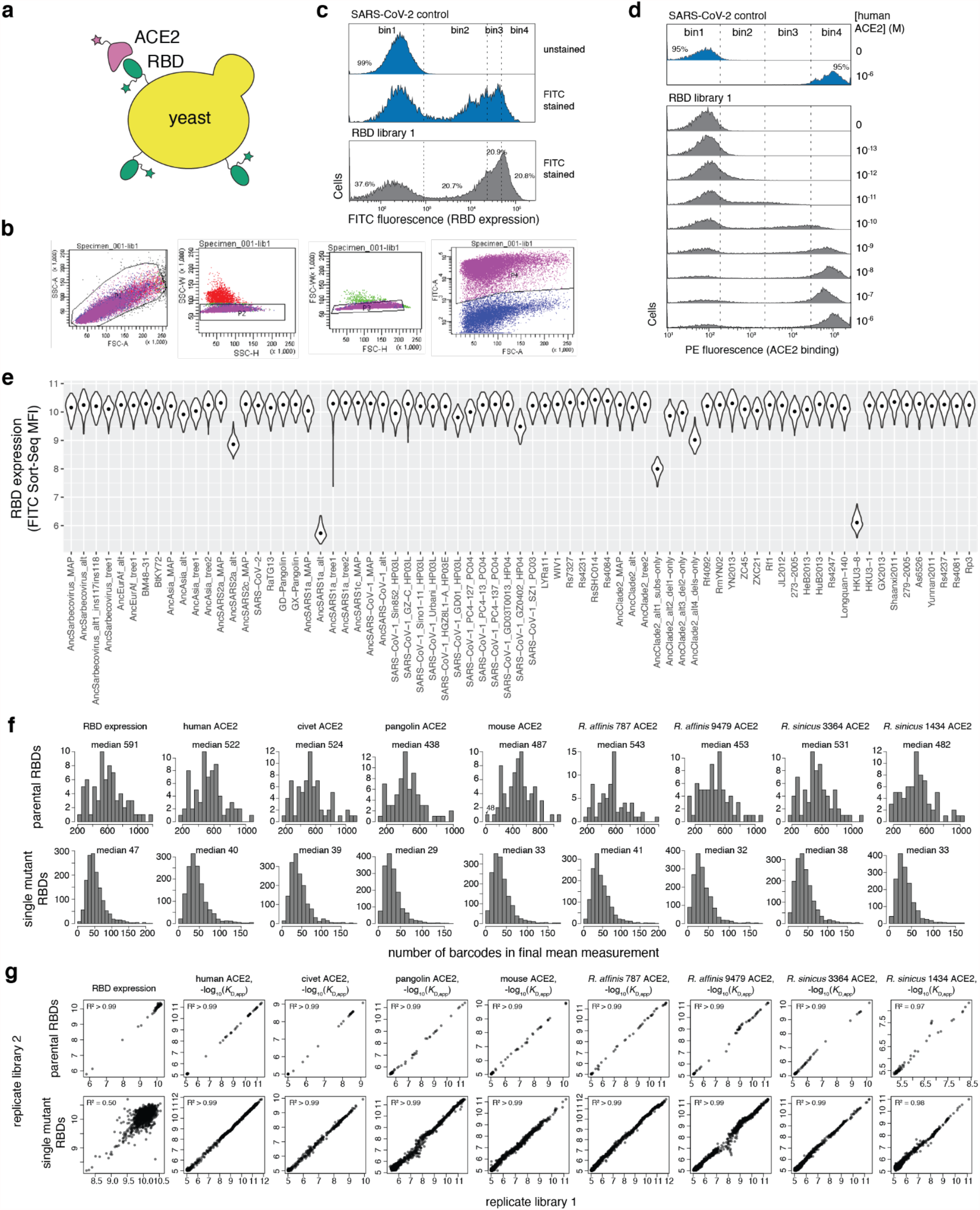
Experimental details of Sort-seq assays. **a**, RBD yeast-surface display enables detection of folded RBD expression and ACE2 binding. **b**, Representative gating for single (SSC-A versus FSC-A, SSC-W versus SSC-H, and FSC-W versus FSC-W), RBD+ (FITC versus FSC-A) cells. **c**, Representative bins drawn on single cells (**b**) for expression Sort-seq measurements. **d**, Representative bins drawn on single, RBD+ (**b**) cells for ACE2 Tite-seq^12,57^ measurements. **e**, Per-variant expression, shown as violin plots across replicate barcodes representing each variant within the gene libraries. **f**, Number of distinct barcodes for each parental (top) or mutant (bottom) RBD genotype used in the determination of final pooled measurements across libraries. **g**, Correlation in measured phenotypes between independently synthesized and barcoded gene library duplicates for parental (top) or mutant (bottom) RBD genotypes.

**Extended Data Figure 3.**
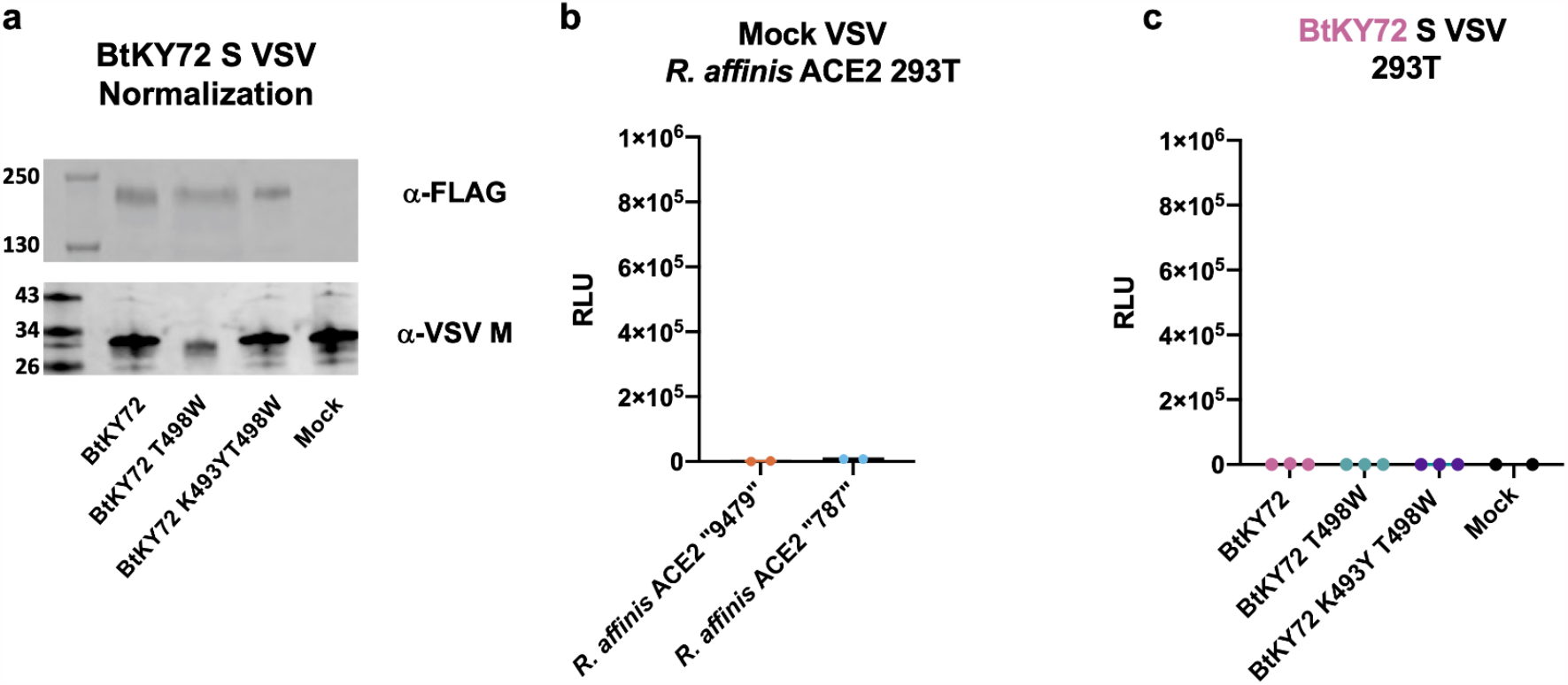
Normalization and controls for pseudovirus entry assays. **a**, Representative Western blots for quantification of spike incorporation into pseudoviral particles. Anti-FLAG identifies incorporation of 3XFLAG-tagged spike, and anti-VSV-M identifies level of VSV backbone. Viral inputs into cell entry assays were normalized across pseudoviral particles by S incorporation as determined in the anti-FLAG Western blot. **b**, Entry into *R. affinis* ACE2-expressing 293T cells by mock VSV particles produced in cells in which no spike gene was transfected. **c**, Entry of pseudoviral particles into 293T cells not transduced to express ACE2.

**Extended Data Figure 4.**
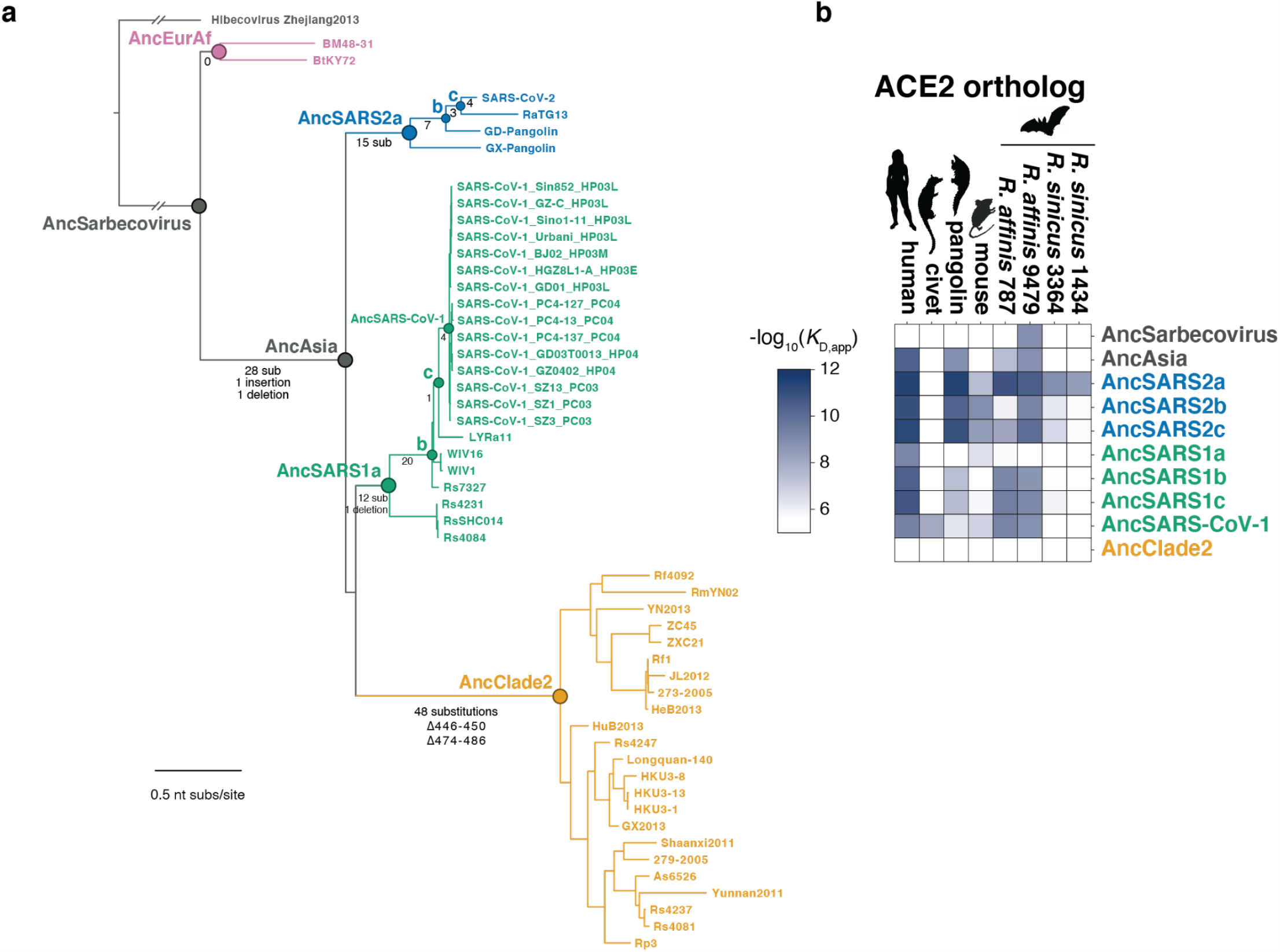
Full set of RBD ancestral sequence reconstructions. **a**, Phylogeny with labeled nodes representing all ancestors tested, including nodes within the SARS-CoV-1 and SARS-CoV-2 clades leading to the human viruses. Branches are annotated with the number of amino-acid substitutions and indels that are inferred to have occurred along each branch. **b**, Phenotypes of all most plausible ancestral sequences (including repetition of the data represented in **Fig. 2b**).

**Extended Data Figure 5.**
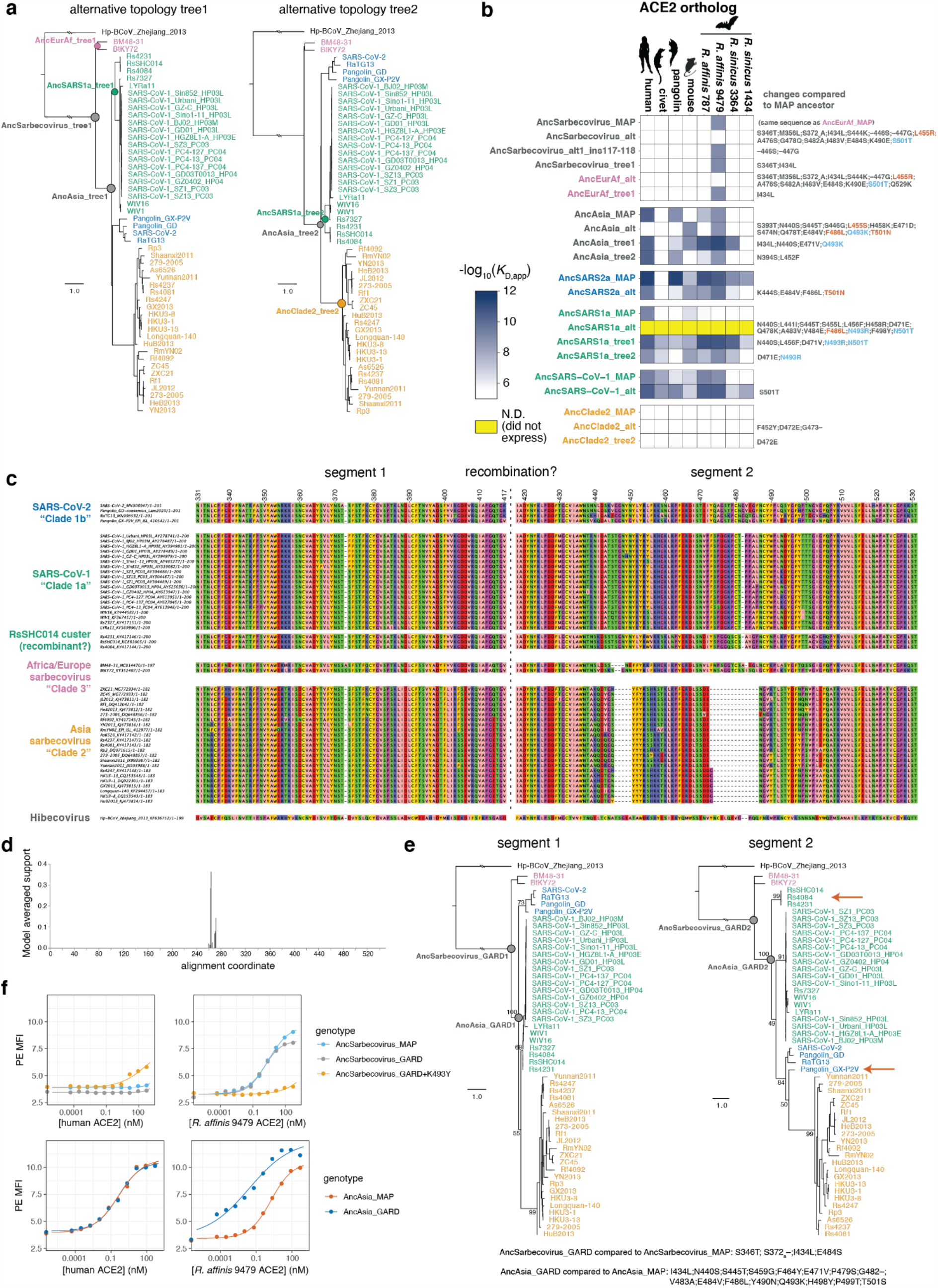
Robustness of ancestral inferences to phylogenetic and statistical uncertainties. **a**, Because of ambiguity in the relationship among the SARS-CoV-1, SARS-CoV-2 and “Clade 2” lineages (**Fig. 1a** and **Extended Data Fig. 1**), we constructed phylogenies constraining a sister relationship between the SARS-CoV-2 clade and Clade 2 (tree1) or the SARS-CoV-1 and SARS-CoV-2 clades (tree2). We performed ancestral sequence reconstruction on these alternative trees. Nodes whose reconstructed sequences differ from the maximum a posteriori (MAP) ancestors shown in **Fig. 2b** and **Extended Data Fig. 4b** are labeled on each tree. **b**, We tested binding of alternative ancestral reconstructions alongside the MAP ancestors in yeast display high-throughput titrations. These alternatives include the “alt” ancestor, which simultaneously incorporates all secondary reconstructed states with posterior probability >0.2^36^; “tree1” and “tree2” ancestors based on the alternative tree topologies shown in (**a**); and for AncSarbecovirus, an alternative “ins117-118” which tests just the ambiguity surrounding the presence or absence of a two-amino-acid indel separate from the remaining amino-acid substitutions tested in AncSarbecovirus_alt. Sequence changes relative to the MAP ancestor are listed at right for each alternative ancestor. For mutations that were tested individually within a background in the mutagenesis data (**Extended Data Fig. 6**), mutations are colored red if they were sufficient to abolish the ancestral phenotype and blue if they reinforced it. Dramatic changes to inferred ancestral phenotypes (and underlying sequence changes in red) are mostly observed in the “alt” ancestors which are the most probabilistically distant test of robustness of ancestral phenotypes, while the tree1 and tree2 alternatives generally recapitulated the MAP phenotypes. The exception is AncSARS1a, where new ACE2 binding capabilities are inferred in the tree1 and tree2 alternatives, but these tree1 and tree2 phenotypes better match what would be expected based on the descendent RBD phenotypes (**Fig. 1b**), suggesting the problem is within the AncSARS1a_MAP reconstructed sequence itself (perhaps due to the presence of recombination, see **c**-**e**). **c**, RBD alignment (shown as amino acids for clarity, though phylogenetic and recombination analysis was inferred on underlying codon nucleotide sequences), with a potential recombination breakpoint identified by GARD^50^ indicated with the dashed line. **d**, GARD relative support values for possible recombination breakpoints. **e**, Phylogenies inferred for the putative non-recombinant RBD segments. Details as in **Fig. 1a**. Arrows point to key changes in the segment 2 sub-tree. The primary recombination signal suggests that the RsSHC014 cluster of RBDs are an independently diverging lineage in segment 2 (which, notably, encodes most ACE2-contact residues), though the placement of GX-Pangolin also shifts in segment 2. We note that each change is supported by only weak bootstrap support values, and this hypothesis introduces a non-parsimonious history with respect to an indel at position 482 in the alignment (**c**). We reconstructed AncSarbecovirus_GARD and AncAsia_GARD on the separate segment 1 and segment 2 trees, and concatenated the matched segments to reconstruct full ancestor representatives accounting for possible recombination. Mutations that distinguish AncAsia_GARD and AncSarbecovirus_GARD from the primary MAP ancestor are listed, bottom. **f**, Key phenotypes of AncSarbecovirus and AncAsia are robust to potential recombination within the RBD alignment. Binding of AncAsia_GARD, AncSarbecovirus_GARD and AncSarbecovirus_GARD+K493Y (see **Extended Data Fig. 6**) to human and *R. affinis* 9479 ACE2 was determined in isogenic yeast display titrations, and are qualitatively unchanged the MAP ancestors.

**Extended Data Figure 6.**
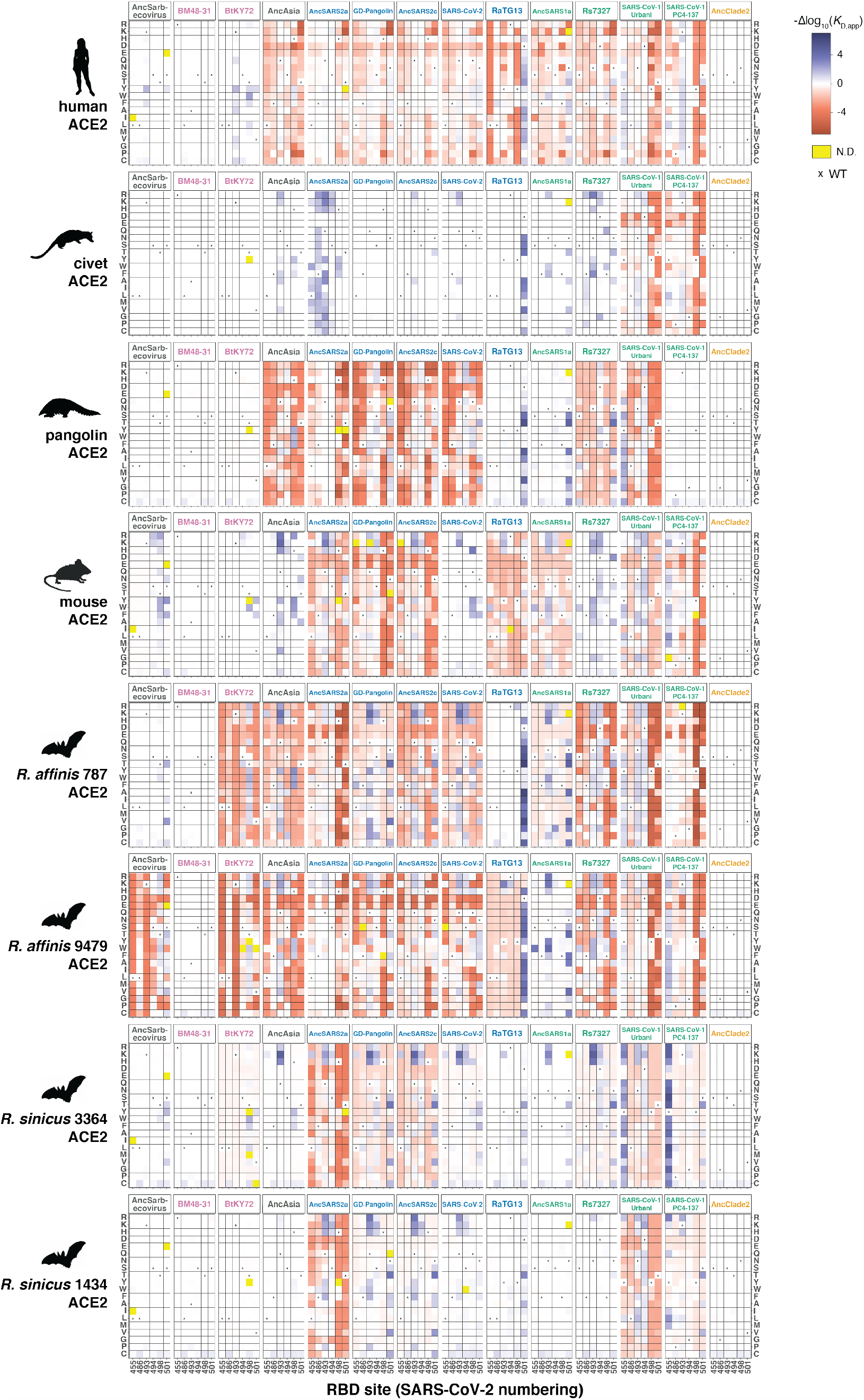
Binding of RBD single mutants to each ACE2. Each heatmap square illustrates the change in binding caused by the indicated mutation at the indicated position (SARS-CoV-2 numbering), according to the scale bar at upper-right. Yellow, mutations that were absent from the library or not sampled with sufficient depth in a particular experiment. x markers indicate the wildtype state at each position in each RBD background.

**Extended Data Figure 7.**
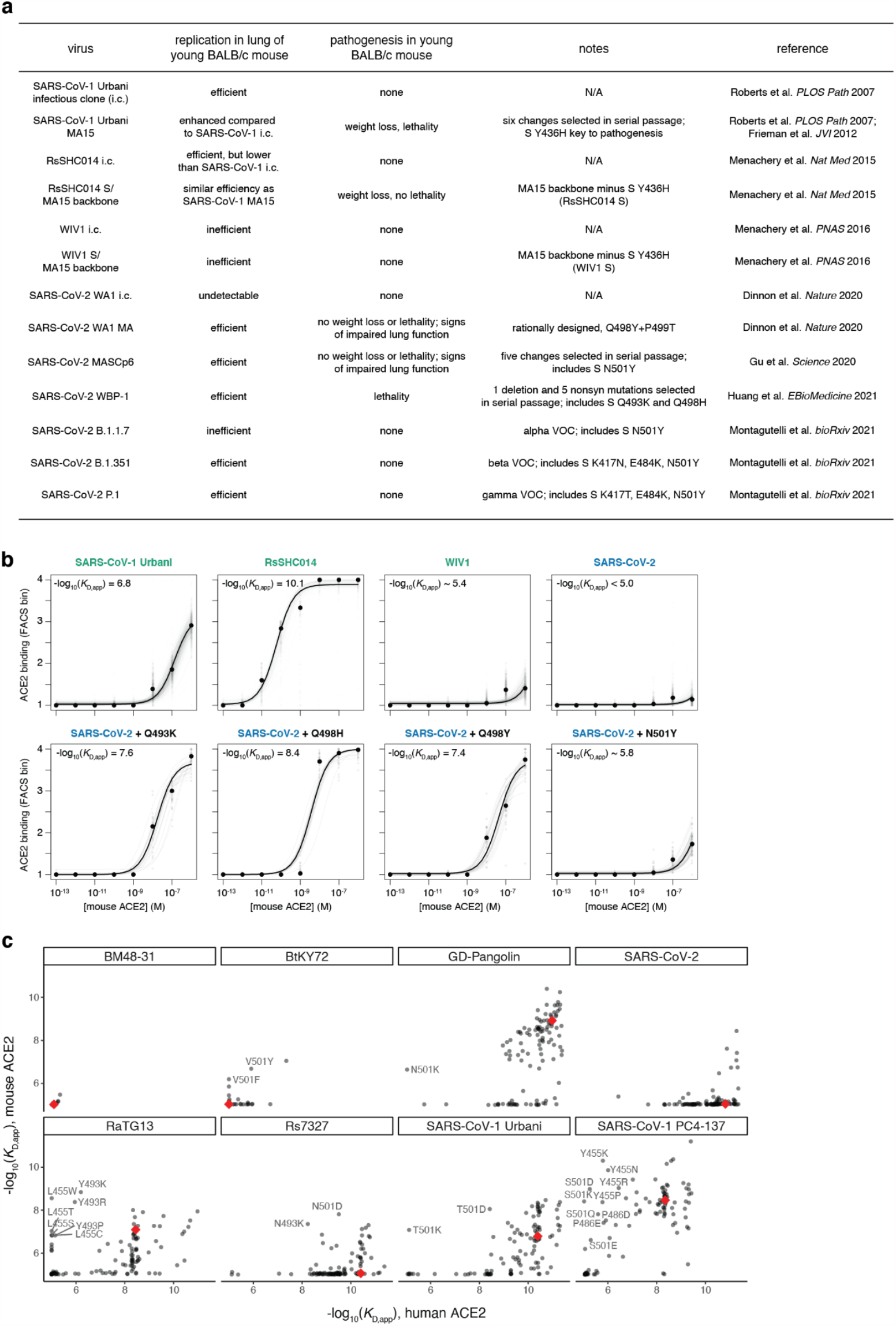
Existing data on sarbecoviruses in mice, and affinities of RBDs and key mutants for mouse versus human ACE2. **a**, Summary of infectivity and pathogenesis of natural sarbecovirus and mouse-adapted strains from prior studies^19,20,40,41,58–61^. **b**, High-throughput titration curves for relevant genotypes from (**a**). Details as in **Fig. 1d**. Strength of binding to mouse ACE2 explains the infectivity and pathogenesis of SARS-CoV-1 Urbani and RsSHC014^19,41^, relative to the weak or absent replication of WIV1^20^ and SARS-CoV-2^40^ in mice. Mutagenesis data explain the inefficient mouse infectivity of the SARS-CoV-2 B.1.1.7 strain^61^ which incorporates the N501Y RBD mutation, relative to the efficient replication of the mouse-adapted SARS-CoV-2 strain containing Q498Y^40^ or the pathogenic WBP-1 strain containing Q493K and Q498H^60^. **c**, An ideal mouse-adapted sarbecovirus strain would bind mouse ACE2 but not human ACE2 for biosafety considerations. The large red points indicate the affinity of the parental RBD for human and mouse ACE2. The smaller black points indicate mutations, and key mutations that enhance binding to mouse versus human ACE2 are labeled. Further mouse ACE2 specificity may be enabled via mutations at other positions not surveyed in our set of six positions.

**Extended Data Fig. 8.**
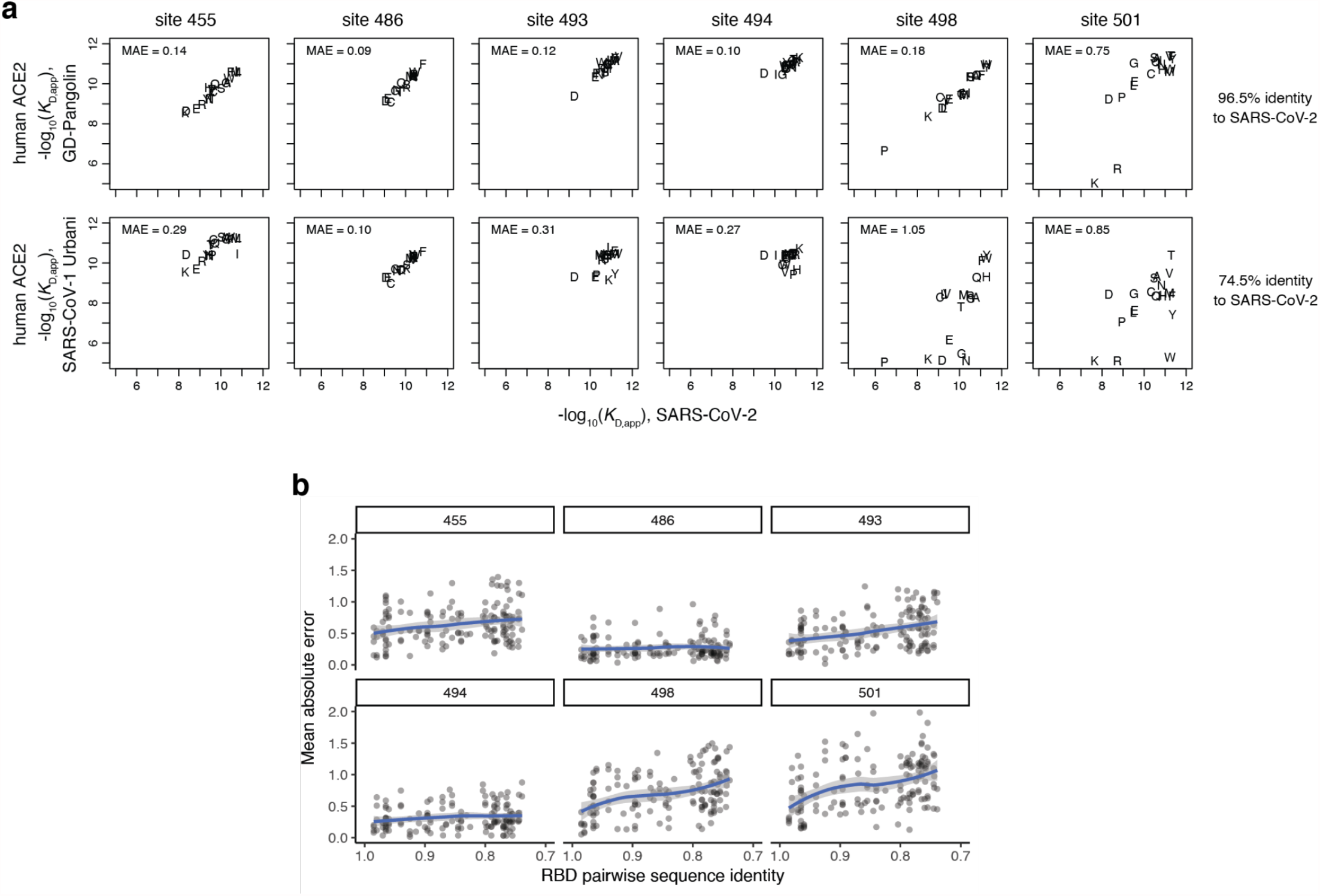
Epistasis and turnover in mutational effects. **a**, Example correlations in binding affinities for mutants at each site for human ACE2. Plots illustrate mutant affinities for human ACE2 and mean absolute error (residual) in the correlation for mutation measurements in GD-Pangolin (top) and SARS-CoV-1 Urbani (bottom) versus SARS-CoV-2. Plotting symbols indicate amino acid for each measurement. **b**, Epistatic turnover in mutational effects across RBD backgrounds. Details as in **Fig. 3g**, but incorporating mutation effects among RBD pairs across all tested ACE2s.

